# Computational modelling of schizophrenia-associated alterations of ion-channel-encoding gene expression predicts a decrease in delta power

**DOI:** 10.1101/2025.10.20.683062

**Authors:** Jan Fredrik Kismul, Torbjørn V. Ness, Torbjørn Elvsåshagen, Ole Andreassen, Gaute T. Einevoll, Marja-Leena Linne, Tuomo Mäki-Marttunen

## Abstract

Schizophrenia presents with a wide range of phenotypes that can help to understand the the mechanisms of the disease. Among these, alterations in delta oscillations are especially amenable to experimental investigation, yet the mechanisms underlying these changes remain insufficiently understood. Biophysically detailed computational modeling offers a powerful approach to investigate these phenomena, as it enables multi-scale integration of genetic and electrophysiological data. In this study, we developed a minimal network model composed of biophysically detailed, multicompartmental neurons to replicate experimental data on the effects of pharmacological blockage of gabaergic neurotransmission on delta-band power. We inserted post-mortem RNA expression data from the anterior cingulate and prefrontal cortices of individuals with schizophrenia and matched controls into the model to study the effects of schizophrenia-associated alterations of ion-channel expression on delta-oscillation power. Our simulations revealed a significant reduction in delta-band power in schizophrenia, driven by altered expression of calcium channel genes in pyramidal neurons. These results provide insights into the genetic contributions to oscillatory disruptions observed in schizophrenia, and our modelling framework can help to develop stratification strategies that bridge genetics and in vivo electrophysiology.

## 1 Introduction

Neuropsychiatric disorders exhibit alterations in brain oscillations in different frequency ranges that can reveal important aspects of the underlying disease mechanisms [1]. This is particularly true for the delta waves (∼0.5–5 Hz), which are associated with unfocused states such as unconsciousness and slow wave sleep [2, 3]. Several psychiatric disorders, schizophrenia (SCZ) in particular, show noticeable alterations in the power of the delta spectrum[4, 5, 6]. SCZ has traditionally been associated with an increased delta power, but observations of a decreased delta power have also been made almost as often [6]. Here, we aim to clarify the mechanisms of the phenotype by simulating the genetic contributions to the alterations of delta oscillations in SCZ through computational modelling.

Delta oscillations are hypothesized to originate either from thalamus or cortex [7]. The cortically generated delta oscillations are likely to depend on the intrinsic properties of layer V pyramidal cells (L5PCs) [7], which is supported by spontaneously emerging delta oscillation-like activity in computational models of L5PCs connected by glutamatergic synapses [8, 5]. Many neurotransmitter systems affect and shape the network activity in the delta band, as shown by the effects of multiple pharmacological agents on delta oscillation power in rat coronal slices [9]. In particular, GABA_B_ receptor activity has been found to be an important factor in the alterations of delta power [9, 10].

Schizophrenia is a highly heritable disorder, with many ion channels implicated [11]. The exact consequences of the genetic variants associated with the risk of schizophrenia are not known, but by combining computational modeling with gene expression data we can study the effects of altered expression [12] at the cellular and network levels, including their contribution to delta oscillations.

Computational models exhibiting delta oscillations exist, but they are either simplified in terms of ion channels [9, 13], or they do not correctly describe the molecular dependencies of neurotransmitters such as GABA_B_-receptor activation [5]. The lack of detail in ion channel descriptions hinders the use of genetics data as there would be no mechanism in which alterations in ion channel functioning could be included. Here we aim to bridge this gap by introducing a layer V cortical network model with pyramidal cells (PC) and two types of interneurons adjusted to fit to data under GABA_B_ receptor blockage.

In this study, we built upon a previous model of cortically generated delta oscillations [5] and added models of basket cells and neurogliaform cells to describe GABA_A_R- and GABA_B_R-mediated inhibition. We connected these interneuron populations to each other and to the recurrently connected L5PC, and we included stochastic glutamatergic inputs to activate the L5PC population. We then fitted the model parameters in order to reproduce data on delta power when blocking GABA_A_ and GABA_B_ receptors separately as measured in ex-vivo electrophysiological experiments [9]. We identified the scizophrenia risk genes encoding ion channels and other proteins included in the model based on a genome-wide association study (GWAS) [14] and altered the corresponding protein densities in our model according to post-mortem expression data [12].

The model predicted that the gene-expression alterations in patients with schizophrenia, particularly those of CACNA1C and CACNA1D, lead to a significant decrease in the average delta power. Our modelling work sheds light on the genetic mechanisms through which delta oscillations are decreased in SCZ patients and paves the way for personalized, biophysics-based approaches in stratification and treatment of patients.

## 2 Methods

### 2.1 Single-neuron Models

We used experimentally validated multi-compartmental neuron models [15, 16] for our modelling analysis. To reduce the computational cost of the simulations, we used simplified-morphology versions of the neuron models as described below.

### 2.1.1 L5PC Neuron Model

A population of thick-tufted Layer 5 Pyramidal Cells (L5PC) were represented by a simplified version of the Hay model [15], as done previously [8]. This model preserved the salient electrophysiological properties of the full scale Hay model while it was 20 times faster to simulate [8].

### 2.1.2 Interneuron Models

The role of GABA_B_ receptors in slow and delta oscillations has been examined in several studies,[17, 18, 19, 20, 21], reviewed in [10]. Large basket cells (LBCs) and GABA_A_R-mediated currents caused by them have been included in a previous network model expressing delta oscillations [5]. While GABA_B_ receptors are involved in many neuron types postsynaptically, it is unclear which neuron types transmit the GABA that activates these receptors. The most likely neurons that mediate this interaction are the neurogliaform cells (NGC). These neurons are one of the few types shown to induce mono-synaptic GABA_B_ currents at a cellular level with a measurable post-synaptic effect [22, 23, 24, 25]. In this work, both basket cell and NGC populations were included, where the basket cells induced GABA_A_R-mediated currents in the postsynaptic cells while the NGCs induced GABA_B_R-mediated currents. The GABA_B_R dynamics were modeled as in the GABA_A_ receptors used in [5], but the rise and decay time constants of the conductance were set to 30 ms and 200 ms respectively, as suggested by experimental data [26].

The single-cell models were selected from the Digital Reconstruction of Neocortical Microcircuit, of the Blue Brain Project (https://bbp.epfl.ch/nmc-portal/microcircuit.html), where we used the `cSTUT189_L5_LBC_29803173bd` model for the basket cells, and the NGC neuron is described by the `L5_NGC_bNAC219_1` model. These models are relatively complex, consisting of 153 and 151 compartments respectively, see Figure 1 panels A and C, making large scale simulations computationally expensive. In order to simplify these models we used the following scheme.

**Figure 1.**
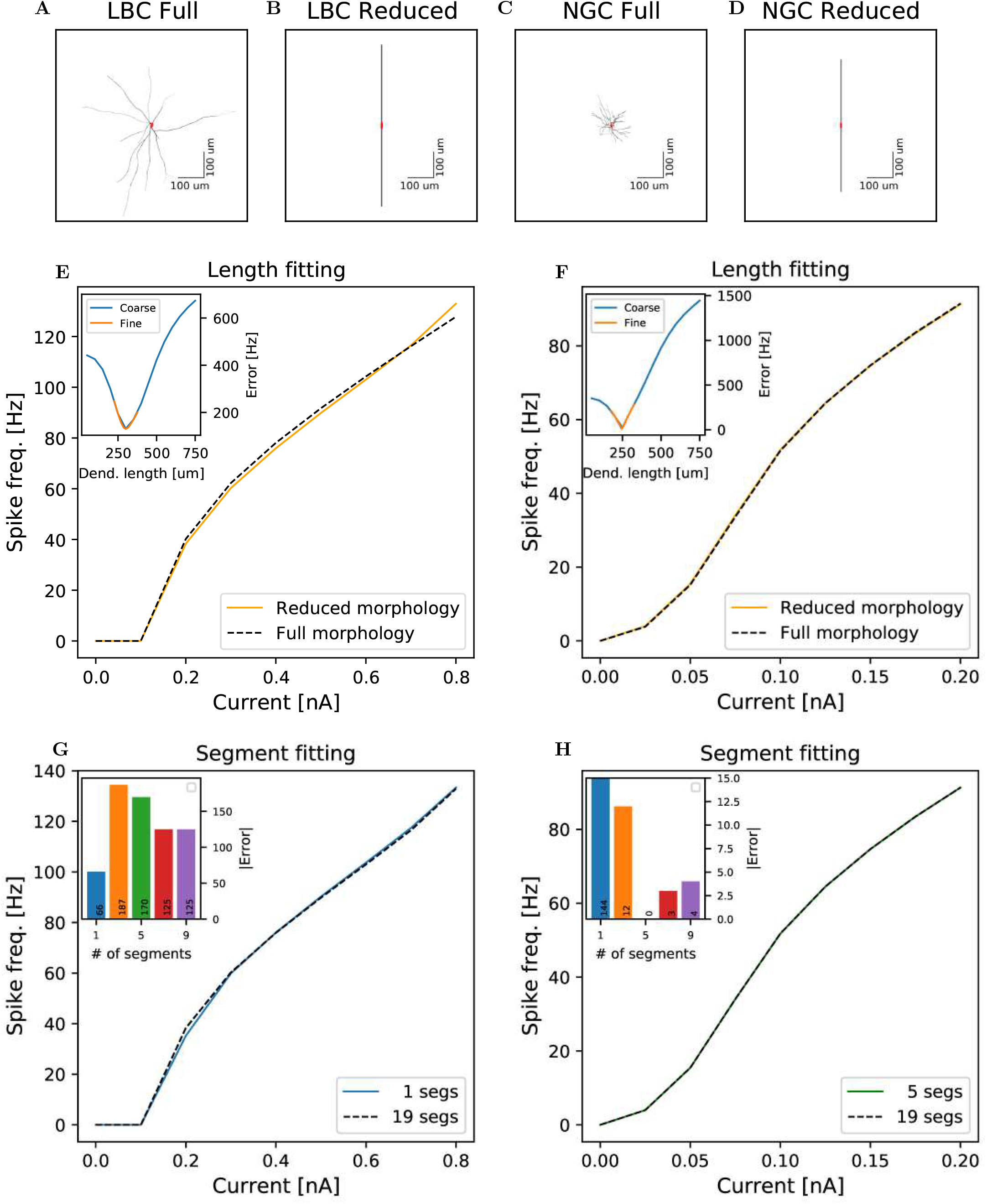
Fitting of reduced-morphology interneuron models. **A-D:** Complete and reduced morhologies of LBC(**A-B**) and NGC(**C-D**) neurons respectively. **E-F:** f-I curves of the complete morphology model versus the best-fit reduced-morphology models for LBC(**E**) and NGC(**F**). Inset: Absolute error of various reduced morphology dendrite lengths; in blue a grid search over a large range of lengths, with a large step length; in orange a fitting around the lowest error range with smaller step lengths to zoom in on the best fit. **G-H:** A fitting of various numbers of compartments per dendrite in the reduced morphologies of LBC(**G**) and NGC(**H**), where 19 compartments per dendrite was the value used when fitting lengths. Inset: Error values for 1, 3, 5, 7 and 9 compartments.

We used ball-and-stick models of 2 dendritic compartments (Figure 1B,D), where each initially consisted of 19 subcompartments. We varied the dendrite lengths, adjusting the dendrite diameter so that we conserved the total dendritic area, in order to replicate the f-I (frequency-current)-curves of the full-morphology models. In this fitting the cell soma received constant direct current stimuli for 20 ms, of 0–0.8 nA and 0–0.2 nA in steps of 0.1 nA and 0.025 nA for LBC and NGC respectively. Currents larger than these led to unrealistically high firing frequencies (*>*100 Hz.) We used a short delay of 200 ms before the onset of the stimuli to allow the neuron to reach an equilibrium state. We then chose the dendrite lengths that minimized the difference to the firing frequencies predicted by the full model.

We first used a coarse grid of dendritic lengths, from 50 - 750 *µ*m in steps of 50 *µ*m, to locate any minima of the error between the f-I curves (i.e., the firing frequencies with respect to the input current amplitude) between full- and reduced-morphology models. We then repeated the grid search using a finer grid around this minima, *±*75 *µ*m in either direction, in steps of 1 *µ*m. The length with the lowest error (301 *µ*m for LBC, 245 *µ*m for NGC) was set as the default length, see Figure 1E,F. Next, we aimed to further decrease the computational complexity of the reduced-morphology interneurons. We now varied the number of subcompartments per dendrite through small odd integers, and compared the firing rates to find the model with the fewest possible segments that still yielded results in accordance with the 19 segment model, see Figure 1G,H. We found best correspondence to the 19 segment models with 1 segment per dendrite for the LBC neurons and 5 segments per dendrite for the NGC neurons.

All mechanisms were kept as in the original full scale models, except for StochKV, which was substituted for a deterministic model developed by Michael Hines https://github.com/OpenSourceBrain/BlueBrainProjectShowcase/tree/master/NMC/NEURON/test/StochKv_deterministic.mod. This was done to permit running simulations with adaptive timesteps, allowing for faster runs.

### 2.2 Network

Our network consisted of 120 PCs, 30 LBCs and 30 NGCs. We connected the neurons randomly with each other in accordance with connection probabilities and numbers of synapses [16]. The network was driven by stochastic input into the AMPA, NMDA and GABA receptors of the PC population only. As the total number of cells in this network is significantly lower than the number of cells in [16] (Figure 2A), we used 6 times larger connection probabilities to minimize the number of non-connected neurons in the network.

**Figure 2.**
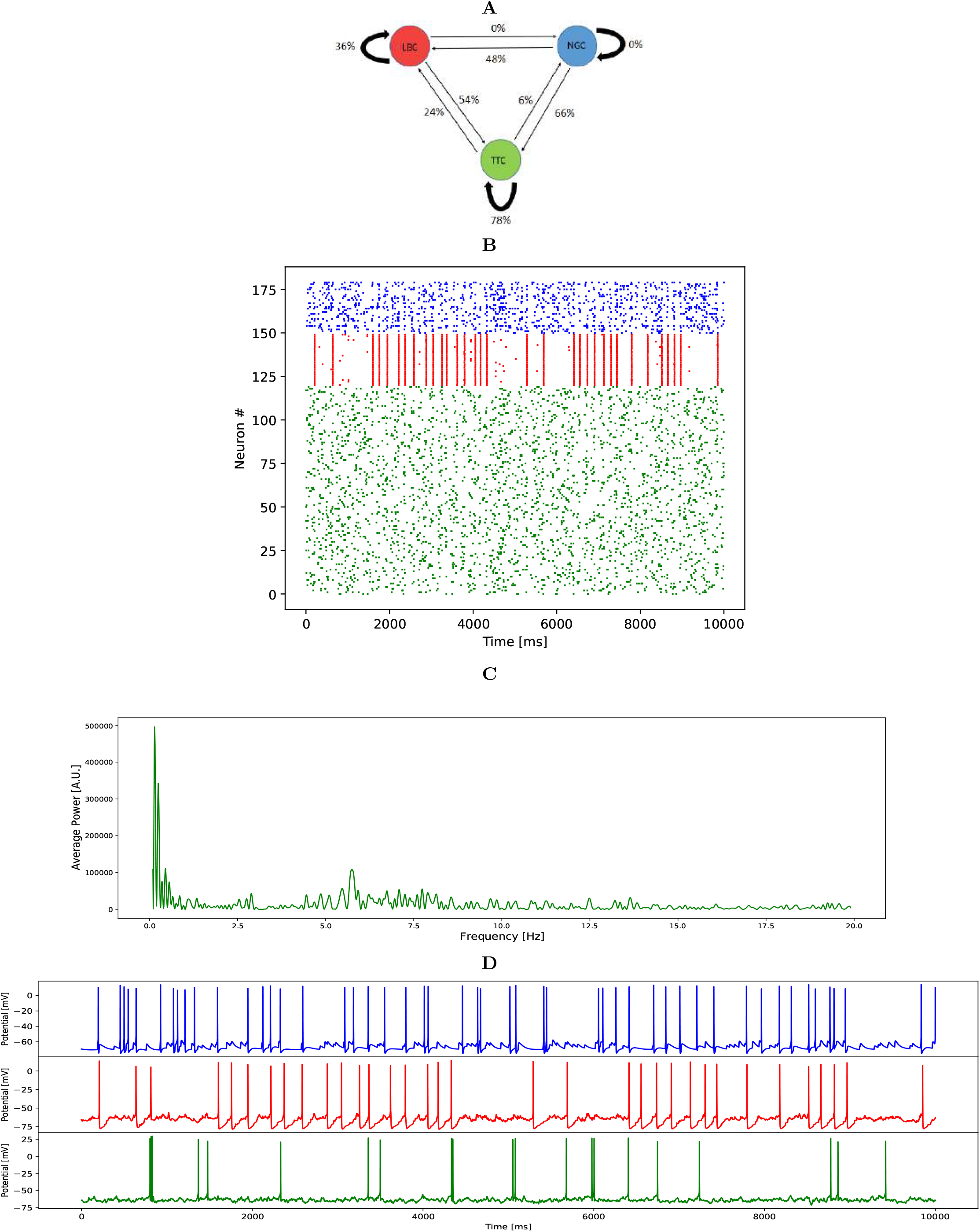
Illustration of the connections, firing profiles and the spectral power of the model network. **A**: Connection diagram between populations, with connection probabilities derived from the literature [16] and scaled by a factor of 6. **B**: Population spike trains of all neurons in the control network model. **C**: Sample power spectrum for the PC population from 0 Hz to 20 Hz. **D**: Soma potentials of a single neuron from each population; NGC(blue), LBC(red) and PC(green).

To make sure the model behaves realistically under altered GABAergic neurotransmission we fit synaptic conductances to data from an earlier study [9], which found that blocking GABA_A_-receptor-mediated currents via gabazine increased the mean delta power to 109% of control, while blocking GABA_B_-receptor-mediated currents via CGP55845 reduced the mean delta power to 17% of control. In our study, this blocking was implemented by setting the conductance of connections from LBC to PC to zero (GABA_A_), or the conductances of connections from NGC to either of the other populations to zero (GABA_B_).

We performed a grid search to fit the synaptic conductances of the network model to the above pharmacological data. Apart from the synaptic connections for which a zero connection proability was reported [16], we varied conductances of all synaptic connections, namely, PC→ PC, PC→LBC, PC→NGC, LBC→PC, LBC→LBC, NGC→LBC, and NGC→PC. To reduce the dimensions of the grid search, we set the conductance from NGC to PC equal to that from NGC to LBC and the conductance from LBC to PC equal to the one from LBC to LBC. This left us with the conductances g_*EE*_ (PC→PC), g_*EI*_ (PC→LBC and NGC), g_*EN*_ (PC→NGC), g_*I*_ (LBC→PC and NGC) and g_*N*_ (NGC→PC and LBC) to vary. Our optimisation task consists of two error functions to optimize, one to capture the GABA_A_ receptor blockage-induced increase in delta power and the other one to capture that induced by GABA_B_ receptor blockage. In addition we ruled out networks with unrealistic spontaneous firing rates.

The resting-state firing rate of the PC population has been reported to be in the range from 1 Hz to 6 Hz in the literature [27, 28, 29, 30, 31], and we chose to aim at a PC firing rate of 3.0 Hz as in our earlier paper [5] (firing rates in the interval [2.5–3.5] Hz were accepted). For the LBC population spontaneous firing rates ranging from 2 Hz to 12 Hz have been reported, namely, 2.1 Hz or 3.0 Hz in mouse visual cortex, depending on the optogenetic tool [32], 3.08 Hz in irradiated rats and 4.89 Hz in healthy rats in dysplastic cortex [33], 5.5 Hz [34] or 9.4 Hz [35] in mouse auditory cortex, and 12.4 Hz in mouse entorhinal cortex [36]. Here, firing rates in the interval [3.0–12.0] Hz were accepted.

We first fitted the conductances by hand to obtain a model that had a qualitatively correct behaviour. We then performed the grid search varying each of the parameters by *±* 7–20% as follows: g_*EE*_ = [0.2,0.25,0.3], g_*I*_ = [0.5,0.6,0.7], g_*EI*_ = [0.7,0.75,0.8], g_*EN*_ = [1.2,1.4,1.6], g_*N*_ = [0.6,0.8,1.0]. For each error function evaluation, we used N=3 repetitions with different random number seeds. After the grid search, we used N=5 repetitions for the parameter sets that fulfilled the firing-rate conditions — these conditions resulted in 3 acceptable networks. In order to select one we calculated the absolute difference between the network alterations in delta power resulting from blocking each of the GABA receptors, and selected the network with the smallest difference. The best fit with our metric was found with g_*EE*_=0.2, g_*I*_ =0.5, g_*EI*_ =0.7, g_*EN*_ =1.2, g_*N*_ =0.6. The activity of the resulting network is illustrated in Figure 2B, and its spectral power distribution and individual membrane potential time courses are illustrated in Figure 2C-D.

### 2.3 Ion-channel gene expression data to inform the model

To model the difference between network activity in healthy controls and SCZ patients, we used the gene expression data from a multi-cohort study of postmortem brains from controls and individuals with SCZ [12].

In this work, we considered gene-expression data from those genes that were implicated as risk genes in [14], i.e., genes where the minimal p-value for a single SNP was smaller than 5 *×* 10^*−*6^. Of the genes whose products were described in our neuron models, six ion-channel-subunit encoding genes and one GABA_B_R-subunit encoding gene met this threshold (listed in Table 1). We used RNA expression sequencing data, normalized by DEseq2 [37], from two cortical brain regions, ACC (Anterior Cingulate Cortex) and PFC (PreFrontal Cortex), extracted post-mortem from 481 (230 SCZ, 251 HC) and 558 (263 SCZ, 295 HC) subjects, respectively. We calculated the average difference in the expression of the seven genes between SCZ and HC populations, see Table 1. In the SCZ models, the conductances of the ion channels encoded by the listed genes were multiplied by the expression ratio (SCZ vs. HC) of these genes. The sole exception was CACNA1C and CACNA1D, which both affect HVA Ca^2+^ ion channel, and therefore the average of the expression ratios of the two genes two was used as a coefficient for this ion channel. Although the gene-expression coefficients of Table 1 were larger than 1 for all considered genes, this was not a consistent trend genome-wide. Namely, out of the 37049 genes whose expression was successfully imputed from the ACC dataset (31681 in PFC), 16007 (43%) had a decreased expression (gene-expression coefficient smaller than 1) in SCZ (12757, i.e. 40%, in PFC). Even within the 687 neurotransmission-associated genes (see Table S2 in [38]), 44% of the genes that were successfully imputed had a decreased expression in SCZ in the ACC dataset and 42 in PFC.

**Table 1:** Model parameter alterations related to gene-expression changes. The second column shows the modeled ion channel affected by the genetic alteration, and the third and fourth columns show the coefficients for the model parameters for ACC and PFC respectively. CACNA1C and CACNA1D affect the same current species, and the average of the two (ACC: 1.164, PFC: 1.137) is used in simulations. The fifth column shows the minimal p-value of the risk SNPs in each gene according to the schizophrenia GWAS [14].

| Gene | Conductance affected | ACC coefficient | PFC coefficient | p-value |
| --- | --- | --- | --- | --- |
| CACNA1C | HVA Ca<sup>2+</sup> | 1.171 | 1.139 | $1.279 \cdot 10^{-21}$ |
| CACNA1D | HVA Ca<sup>2+</sup> | 1.158 | 1.135 | $3.277 \cdot 10^{-9}$ |
| CACNA1I | LVA Ca<sup>2+</sup> | 1.173 | 1.144 | $1.171 \cdot 10^{-13}$ |
| HCN1 | HCN K<sup>+</sup> | 1.034 | 1.117 | $2.873 \cdot 10^{-14}$ |
| KCNB1 | Persistent K<sup>+</sup> | 1.123 | 1.150 | $2.197 \cdot 10^{-10}$ |
| KCNQ3 | M-type K<sup>+</sup> | 1.056 | 1.034 | $2.128 \cdot 10^{-6}$ |
| GABBR2 | GABA<sub>B</sub>R mediated inhibitory conductance | 1.084 | 1.095 | $9.334 \cdot 10^{-9}$ |

All simulations were performed using the NEURON simulator. Our model scripts are publicly available at ModelDB (https://modeldb.science/2020338, password: delta)

## 3 Results

We simulated a network consisting of 120 PCs, 30 LBCs and 30 NGCs, all interconnected according to Figure 2, and driven by a random background input to the PCs. Recent GWAS [14] and gene-expression data [12], normalized by DEseq2 [37], were used to investigate the effect of the different ion-channel encoding gene variants on the delta power. The effect of the expression alterations as observed in post-mortem PFC or ACC on delta power was tested using population-averaged and subject-wise manipulation of the corresponding ion channel conductances. After this, we analyzed in more detail the effects of these data-driven manipulations in isolation (one ion-channel conductance at a time), and we also tested the contributions of different neuron populations to these alterations by implementing the manipulations in one population at a time.

### 3.1 Alterations of expression of ion-channel-encoding genes in SCZ lead to decreased excitability and altered spectral power

An earlier study [5] analyzed the effects of ion-channel variants of SCZ-associated genes on delta power. Here, we extended this model with realistic GABAergic neuron models and post-mortem expression data from SCZ vs HCs (see Methods) to analyze the effects of the actual expression-level differences of these genes on the predicted delta power.

Simulations of ACC and PFC were first carried out for N=30 repetitions with different random number seeds for the population-averaged expression-level differences in post-mortem PFC and ACC of SCZ patients (see the ion-channel conductance coefficients of Table 1). The predicted delta power was decreased in SCZ in both regions; ACC-like variants caused a 14.2% decrease (U-test; p-value 1.84*·*10^*−*4^), and PFC-like variants caused a 12.4% decrease (U-test; p-value 6.73*·*10^*−*4^), see Figure 3A. The average firing activity per pyramidal neuron was significantly decreased both in the ACC and PFC regions, from an average number of 3.14 spikes per second in healthy controls to 2.97 spikes/s in ACC (U-test: p-value 8.48*·*10^*−*9^), and to 3.04 spikes/s in PFC (U-test: p-value 2.23*·*10^*−*6^). The spectral power in other frequency bands was also affected in SCZ — namely the theta power (5–8 Hz) decreased in ACC (U-test: p-value 2.20*·*10^*−*5^) and PFC (U-test: p-value 1.01*·*10^*−*4^), the alpha power (8–12 Hz) decreased in ACC (U-test: p-value 5.27*·*10^*−*6^) and and PFC (U-test: p-value 5.41*·*10^*−*4^), the beta power (12–30 Hz) decreased in ACC (U-test: p-value 2.98*·*10^*−*6^) and PFC (U-test: p-value 4.58*·*10^*−*4^), and the gamma power (30–150 Hz) in ACC (U-test: p-value 4.58*·*10^*−*6^; Figure 3B).

**Figure 3.**
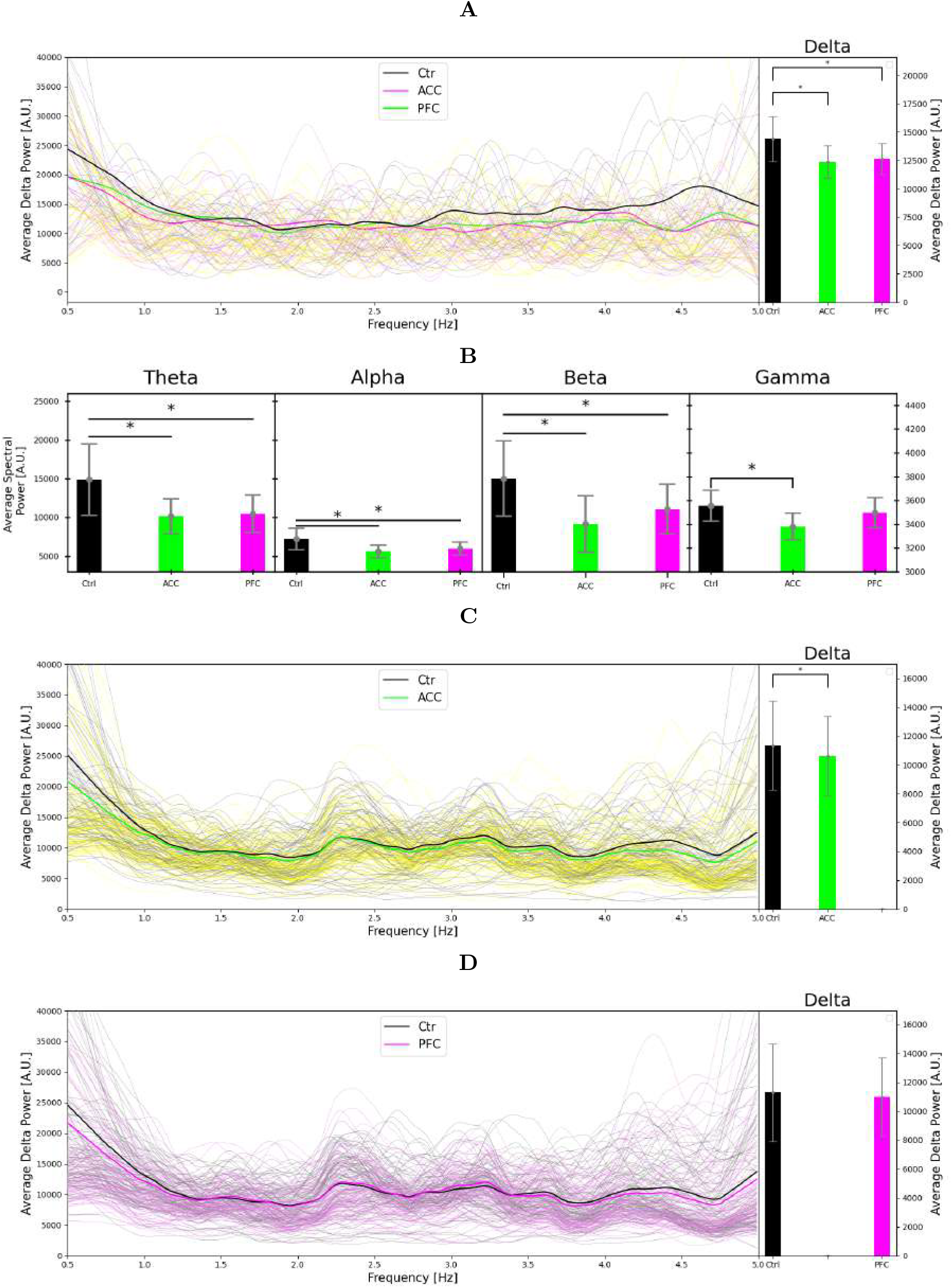
Predicted alterations in delta power based on expression data. **A**: Spectral power (left) and average delta power (right) of L5PC spiking activity from simulations of N=30 networks, smoothed by LOESS [39]. CommonMind data from ACC (purple) and PFC (gray) were used for modelling the effects of population-averaged SCZ-associated alterations of gene expression in the respective brain regions, lighter respective colors representing each random seed. The default network model was used for the HC (brown) case. A significant decrease in delta power was observed for both SCZ groups. **B**: Average spectral power for other frequency bands (theta: 5–8 Hz, alpha: 8–12 Hz, beta: 13–30 Hz, gamma: 30–150 Hz). A significant decrease is seen in the ACC region for all the frequency bands, and in the PFC region for the lower three frequency bands. **C**: Spectral power (left) and average delta power (right) of subject-wise simulations based on ACC gene-expression data. In these simulations, the ion-channel conductance coefficients (see Table 1) were calculated for each subject (both HC and SCZ) by normalizing the individual gene expressions by the mean expression levels in the HC population. The left panel shows the LOESS smoothed simulation results in average in bold colors, and the 100 first subjects in lighter respective colors. The right panel shows the mean and SD of the average delta power across the HC and SCZ populations – a significant decrease in delta power was found for the SCZ group. **D**: The experiment of (B) repeated for PFC. No significant decrease in delta power was found for the SCZ group.

To complement the simulations based on population-averaged gene expression levels, we carried out a subject-wise simulations where the ion-channel conductances were altered by factors representing the relative expression of the considered ion channels in each subject compared to an average HC subject (see Methods). As expected, the standard deviation of the predicted delta power was larger in both CTR and SCZ in the subject-wise analysis compared to the population-averaged simulations. The delta power was significantly decreased in ACC in SCZ subjects compared to HC subjects: the delta power

was on average 6.3% smaller than in HC subjects (U-test: p-value 4.67*·*10^*−*3^), with an average number of spikes per pyramidal neuron per second decreasing from 3.00 to 2.91 (U-test: p-value 3.09*·*10^*−*3^), while in PFC there was no significance decrease, see Figure 3C–D.

Taken together, our model predicts that delta power is significantly decreased in SCZ in both ACC and PFC, both when modeled with average and with subjectwise conductance coefficients.

### 3.2 Predicted decrease in delta power is mediated by altered calcium channel expression in L5PCs

We carried out simulations to investigate which genetic variants had most effect on the delta power, where only a single ionchannel conductance was manipulated. Running 30 simulations for each genetic variant, we found a statistically significant role only for the HVA Ca^2+^ ion-channel variations. This ion channel is affected by the the altered expression of CACNA1C and CACNA1D, with a decrease of 8.9% compared to healthy subjects (U-test: p-value 0.015) for ACC, and a decrease of 11.6% (U-test: p-value 0.001) for PFC, see Figure 4A.

**Figure 4.**
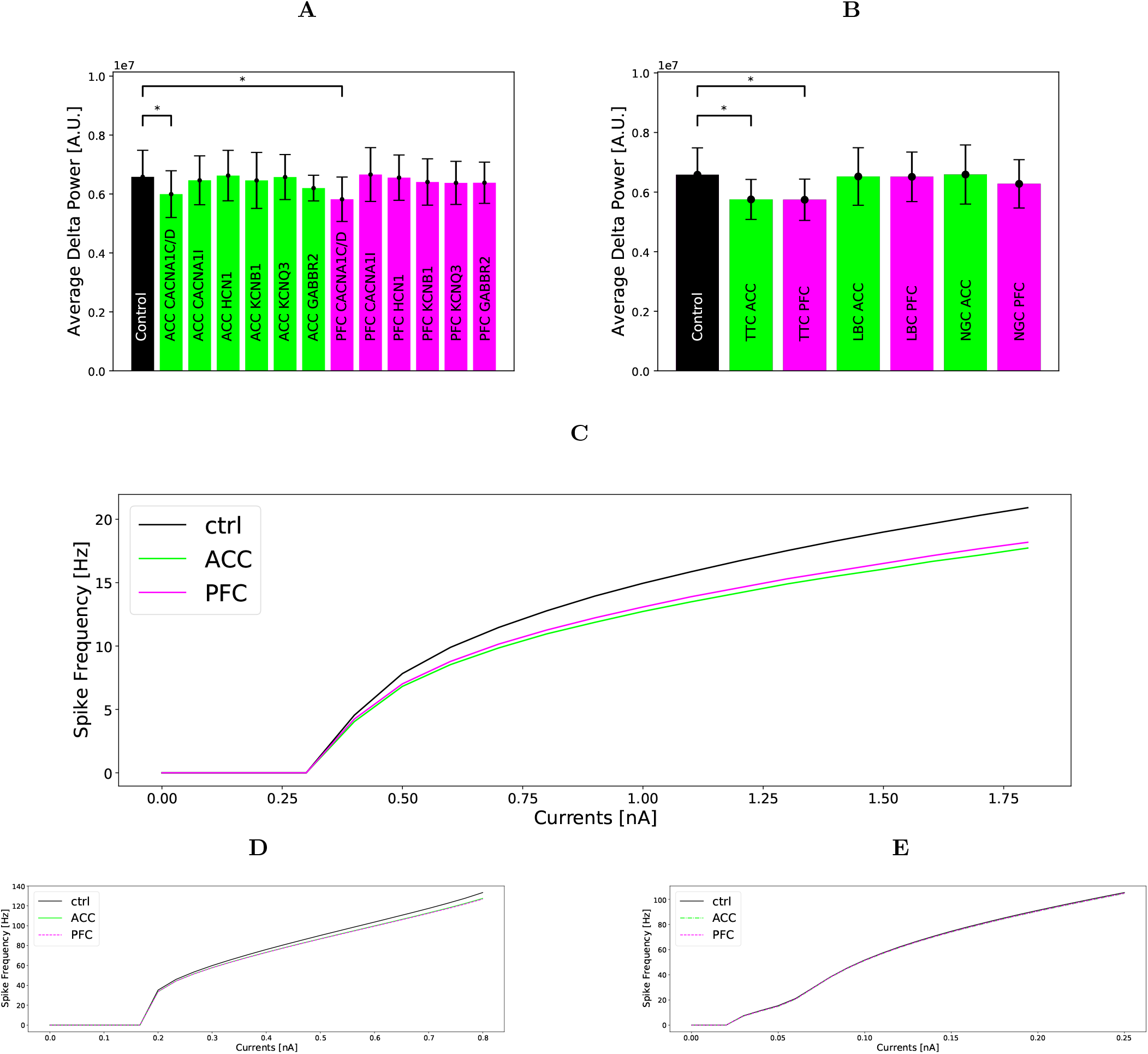
Ion-channel and neuronal population effects on delta power and f-I curves. **A:** Average delta power from simulations of N=30 networks where the SCZ-associated expression-level alterations were implemented in only one neuron population at the time (as opposed to Figure 3 where the ion channel conductances of all neurons were altered). Similar to Figure 3A, we used population-averaged ion-channel coefficients (Table 1). A significant decrease in delta power is found for CACNA1C and CACNA1D for both ACC and PFC. P-values: ACC CACNA1C and CACNA1D 0.015, ACC CACNA1I 0.515, ACC HCN1 0.894, ACC KCNB1 0.626, ACC KCNQ3 0.965, ACC GABBR2 0.450, PFC CACNA1C and CACNA1D 0.001, PFC CACNA1I 0.871, PFC HCN1 0.848, PFC KCNB1 0.408, PFC KCNQ3 0.337, PFC GABBR2 0.762.**B:** Summed delta power from simulations of N=30 random seeds per population for all population, performed with the average population ion-channel coefficients. A significant decrease in delta power is found in the PC population for both ACC and PFC. P-values: ACC PC 6.73·10^*−*4^, LBC 0.723, NGC 0.976, PFC PC 6.37·10^*−*4^, LBC 0.636, NGC 0.183.**C**,**D**,**E:** The effect of genetic variants on the f-I-curves of each simplified neuron; PC(top), LBC(middle), NGC(bottom). Each neuron receives an injected current to the soma for 20 seconds, and the spike frequency is recorded for each injected current.

A population-wise application of the altered ion-channel conductances showed that it was when the variations affected the PC population that the delta power significantly decreased, and that the other two populations seemed to have negligible contributions, see Figure 4B. A U-test of N=30 repetitions of different random seeds showed a 12.6% decrease in delta power (U-test: p-value 6.73*·*10^*−*4^) for ACC, and a decrease of 12.8% in delta power (U-test: p-value 6.37*·*10^*−*4^) for PFC in the PC population.

The stronger effect of genetic variants in the PC population was reflected in the f-I curves for the simplified neurons, shown in Figure 4C-E. The effect of genetic variants in the LBC population is smaller, and the effect on the NGC population was negligible.

Taken together, we find that manipulations of the calcium channels in the pyramidal neuron population have the largest effect on the delta power, while corresponding changes in other cell types as well as manipulation of other parameters according to the gene-expression data have little effect.

## 4 Discussion

In the present work we combined psychiatric post-mortem RNA expression data with computational modelling and found that the SCZ-associated gene-expression alterations predicted a decrease in delta oscillations and that this change is driven primarily by the increased calcium channel expression in PCs (Figure 4A-B). Since this model allows manipulating the conductances of many of the ion channels encoded by SCZ-associated genes, it can be used for the study of other phenotypes of SCZ whose neural mechanisms are sufficiently understood.

Our results for SCZ-associated alterations in delta oscillation power are based on changes in bulk expression of ion-channel-encoding genes obtained from post-mortem brains. Although the expression levels are likely to differ between different cell types in the considered brain areas, qualitatively similar expression-level results were obtained for the most important genes CACNA1C and CACNA1D in a recent paper [38] (CACNA1C +14.87% and +3.73% vs our +17.1% and +13.9% in ACC and PFC, respectively; CACNA1D +8.29% and +1.65% vs our +15.8% and +13.5% in ACC and PFC, respectively) where expression data imputed for neurons were used and corrected for differences in sex, age and post-mortem index. Ultimately, protein-expression measurements should be carried out to further improve the predictive power of our SCZ vs HC modelling. It should be noted that the post-mortem expression data may be affected by the use of antipsychotics. Here, we deliberately left out expression data of glutamatergic and GABA_A_ receptors, since the expression of these proteins is likely to be affected by antipsychotics use [40, 41, 42, 43].

The current network model of layer V was developed taking into account the effects of blocking GABA_A_ and GABA_B_ receptors on the delta power. Beause it describes the GABA_B_-receptor-dependency of the network activity, it can be a valuable tool for studying other phenotypes of schizophrenia that involve GABA_B_-receptor activity. We found that the predicted decrease in delta power observed in SCZ subjects is mostly caused by altered expression of the ion channels expressed in the PC population, an observation that is reinforced by the single neuron f-I curves (Figure 4C-E). This could be partly due to the fact that we based our spectral power estimates on L5PC spiking only. Calculating a spectral power estimate from firing rates (as done in [44]) does not take into account the synaptic activity in the dendrites underlying LFP/EEG signals, however, studies show that such approaches can still reasonably accurately reflect the low-frequency range of LFP/EEG power spectra that we are interested in here [45, 46]. In addition to the ion-channel encoding genes associated with SCZ, we also tested the effects of altered expression of GABBR2, but the delta-oscillation power was not significantly affected by this.

Of the six gene variants affecting relevant ion channels we found only the two affecting the high-voltage activated calcium channel (CACNA1C and CACNA1D) to have a significant effect. We could not distinguish between the effects of these two genes as our model contains only one high voltage activated calcium channel.

There exist many previous network models that describe slow oscillations and delta oscillations, and at least two such models also included a description of GABA_B_R-mediated inhibitory currents [9, 13]. In one of these [13] the neurons are represented by simple integrate and fire neurons. Other differences of note are the difference in number of neurons (4000 vs our 180), amount of inhibitory neurons (17% vs. our 33%), as well as a non-uniform connection scheme. One important distinction is that each of our neuron populations contain neuron-type specific parameter values (morphology, ion-channel conductances and passive parameters) while [13] used the same values for all neuron types. The other model [9], which was based on an earlier model [47], consisted of a model containing more details in several respects; their neuron morphologies were more detailed, the populations were split into two layers, they had 13 populations compared to our 3, and their GABA_B_ receptors had a time delay before spiking. In their model there were gap junctions within each neuron population, while we only have them between LBCs. In addition, our model runs for a much longer runtime. The advantage of our approach is that the neuron models we used were morphologically detailed and their ion-conductances were fitted to single-cell electrophysiological data. This permits more intricate studies where the effect of SCZ-related alterations in ion-channel expression on network activity can be examined.

The predicted decrease in the overall delta power is in accordance with some of the experimental research, while other reports suggest an increase rather than a decrease in the delta-oscillation power [6]. Our results sheds light on these inconsistencies by providing a mechanistic explanation for how differences in the expression of ion-channel-encoding genes can lead to a decreased delta power. The observations of increased delta power could thus be caused by either environmental factors or genetic factors affecting expression of genes other than those encoding ion channels. However, it should be noted that we only considered the effects of altered expression of ion channels in adult cells and networks, while some ion-channel encoding genes, CACNA1C in particular, are known to affect the wiring of a developing brain [48, 49]. Future research should thus be directed toward developing modelling frameworks that permit gene-based modelling of the development of neuronal networks, which would allow predicting how the structure of the cortical network is affected by SCZ-associated alterations in gene expression. Integrating such data into our model would greatly improve the predictive power of our gene-based modelling framework.

Since our model only contains three neuron populations and only describes neurons with their somas in the fifth layer of the cortex, there are limitations on what our model can be used for. However, the model is ideally suited to studying cortically generated delta oscillations since the L5PCs included in the model are in a key position for both delta oscillation generation and the contribution to the EEG signals. Our assumption of the driving signal, the background synaptic firing, being unaffected by the SCZ-associated gene-expression alterations should be reconsidered once there are data and models available that allow realistic manipulation of the background synaptic activity. In particular, incorporation of a thalamic circuit responsible for the thalamically generated delta oscillations [50] and the thalamocortical connections into our model would permit examining the effects of SCZ genetics on the two distinctly originating delta oscillations. Given the link between altered delta oscillations and cognitive symptoms such as working memory deficits [51], our findings and their reproduction in a larger model could lead to new treatments in the future.

Allowing for altering individual ion channel conductances makes the model suited for studying other known phenotypes of SCZ, i.e. P50 suppression or mismatch negativity (MNN). Biophysically detailed modelling of multiple phenotypes in a subject-wise manner, based on genetics data, may help in patient stratification and, in the future, personalized medicine approaches.

